# Bayesian-enhanced closed-loop optimization of ultrasound protocols for targeted and precise neuromodulation

**DOI:** 10.64898/2026.06.16.732762

**Authors:** Andrea Boscutti, Valeria Grasso, Tommaso Di Ianni

## Abstract

Low-intensity focused ultrasound (LIFU) is a promising neuromodulation modality, but challenges related to high response variability and the poorly understood parameter space undermine progress in clinical applications. To facilitate the development of therapeutic LIFU protocols, we developed an approach for Bayesian-enhanced adaptive control of ultrasound neuromodulation (BEACUN). BEACUN enables efficient, data-driven parameter mapping using a limited number of stimulation-response evaluations. We used functional ultrasound imaging (fUSI) to measure the neural responses to LIFU stimulation in real time, and we carried out *in vivo* experiments in rats to optimize and validate the performance of the BEACUN search. In live optimizations, we show that BEACUN produces more effective inhibitory LIFU neuromodulation protocols than conventional parameter exploration methods and converges to the optimal solution in 23 ± 3.67 stimulation-response evaluations. Our approach realizes a platform for efficient optimization of neuromodulation parameters that could pave the way for personalized LIFU protocol development in patients.

## Introduction

Low-intensity focused ultrasound (LIFU) is an emerging noninvasive neuromodulation technique that combines high spatiotemporal resolution with the ability to target deep brain structures^1–3^. Despite its promise, several factors continue to limit broader clinical application of LIFU, chief among which is a poor understanding of how stimulation parameters relate to neural outcomes^4^. Relevant LIFU stimulation parameters include acoustic intensity, pulse repetition frequency (PRF) – i.e., the rate at which ultrasound pulses are delivered – duty cycle (DC) – i.e., the fraction of time over which the stimulation is delivered – and stimulus duration^5,6^. Higher-order features of the stimulation protocol, such as the temporal patterning of pulse trains or the choice of target brain region, further expand the parameter space^7,8^. This challenge is not unique to LIFU, as parameter optimization remains an open problem across non-invasive (e.g., transcranial magnetic stimulation) and invasive (e.g., deep brain stimulation) neuromodulation modalities^9^. However, the broad ensemble of parameter combinations and the possibility of targeting multiple putative therapeutic sites across virtually the entire human brain make parameter mapping in LIFU neuromodulation a particularly onerous endeavor.

Parameter exploration in LIFU neuromodulation research typically relies on grid search, in which a pre-determined, fixed sampling of the parameter space is evaluated. The grid is defined a priori based on existing and often incomplete knowledge of the LIFU stimulation-response function and fixed budget constraints limiting how many stimulation-response evaluations can be performed in each subject. As a result, only a small subset of all possible conditions is tested in typical experimental settings. Furthermore, this approach has a key limitation in that observations are treated as independent, and no information from previous experiments is used to guide subsequent parameter choices. These challenges are particularly restrictive for the development of optimized LIFU protocols in clinical applications, where time and resource constraints, along with substantial neurobiological heterogeneity, make it difficult to carry out an exhaustive parameter search. In addition, the high complexity of the parameter mapping task precludes treatment personalization, which has been shown to be critical to enhancing the safety and therapeutic efficacy of neuromodulation treatments^9–11^.

To address these limitations, here we developed and validated a method for Bayesian-enhanced adaptive control of ultrasound neuromodulation (BEACUN). BEACUN is a machine learning-based approach for effective mapping of LIFU parameters using Bayesian optimization (BO) with Gaussian process (GP) regression. BO enables data-driven parameter mapping by estimating the optimum of an unknown (black-box) objective function using a limited number of stimulation-response evaluations^12,13^. Compared to other optimization methods frequently used in experimental settings, BO converges faster and yields more robust solutions, as demonstrated in varied applications including chemical reaction development^14,15^, robotic chemistry^16^, and optimization of computational imaging approaches^17^. Versions of BO have been used previously for protocol optimization with other neurostimulation modalities including deep brain stimulation^18–22^, transcranial magnetic stimulation^23^, transcranial current stimulation^24^, vagus nerve stimulation^25^, and neuroprosthetics^26–29^.

We validated our BEACUN approach for effective online mapping of LIFU neuromodulation parameters through *in silico* and *in vivo* experiments in a rat model. To this end, we combined LIFU with functional ultrasound imaging (fUSI), a neuroimaging modality that tracks changes in cerebral blood volume (CBV) in the brain microvasculature with high resolution and sensitivity^30–32^. The CBV signal provides an indirect measure of local neural activity via neurovascular coupling^33^. In our experiments, we used the fUSI readout as a measure of the neural responses to the LIFU stimulation. We performed an extensive mapping of the fUSI-LIFU parameter space and built a virtual experimental model based on the ground-truth data to benchmark the performance of the BEACUN approach against conventional optimization methods. We then deployed the BEACUN platform in a live, fully automated LIFU parameter search in the presence of event-related neural activations, demonstrating substantially greater efficiency compared to conventional stimulation mapping methods.

## Results

### Bayesian optimization in LIFU neuromodulation

We define the LIFU parameter space ***X*** as the *n*-dimensional set of independent variables representing the stimulation parameters. The objective function *f*(***X***) is the unknown stimulation-response function, which represents the evolution of a response biomarker (e.g., functional neural activity, network connectivity, behavioral measure) modulated by the LIFU stimulation as a function of the parameters ***X*** (**Figure 1**).

**Figure 1:**
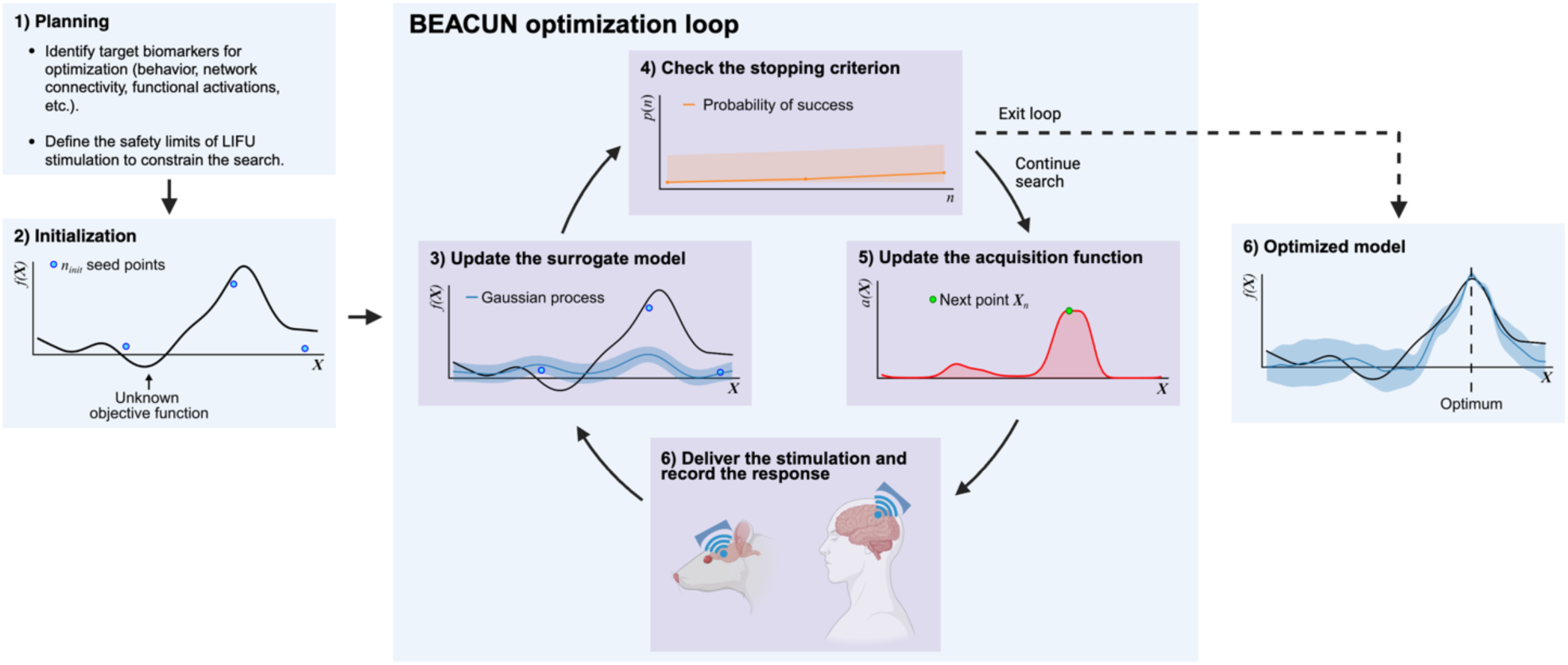
Bayesian-enhanced adaptive control of ultrasound neuromodulation. Schematic representation of the Bayesian optimization loop in low-intensity focused ultrasound neuromodulation.

In the initialization phase, *n_init_* stimulations are carried out with randomly distributed parameter configurations ***X****_init_*. A response *f̅*(***X****_init_*) is recorded after each evaluation, and all the initial responses are used to build the first surrogate model *f̂*(***X***) of the objective function (*prior*) via GP regression. An acquisition function selects the next candidate ***X****_n_* to test (**Supplementary Figure 2b, 2e**), the physical device delivers the LIFU stimulation using the chosen parameter set, and a response *f̅*(***X****_n_*) is recorded (**Figure 1**). The surrogate model *f̂*(***X***) (*posterior*) is then updated via GP regression, and the process is repeated iteratively until a stopping criterion is met. The stopping criterion can be defined based on a fixed, predetermined budget (e.g., the number of evaluations that are deemed feasible in a patient population or the resources allocated to the optimization procedure), or it can be automated using probabilistic measures of success. At the end of the BEACUN loop, the optimum of the GP model *f̂*(***X***) tends to the optimum of the objective function *f*(***X***) (**Figure 1**). To tune our BEACUN implementation in LIFU optimization tasks, we optimized the acquisition function, the number *n*_!"!#_ of initial points, and the initialization strategy.

### fUSI-LIFU ground-truth parameter mapping

To determine the impact of relevant BO hyperparameters and benchmark the performance of our BEACUN approach against a realistic ground-truth objective function, we carried out an extensive parameter mapping in a LIFU neuromodulation application in rats (**Figure 2**). We first characterized the LIFU acoustic field through hydrophone measurements in water (**Figure 2b**). Then, we directed LIFU stimuli to the superior colliculus (SC) unilaterally (**Figure 2a-c**) and interleaved the stimuli with fUSI recordings to measure the neural responses to the stimulation. We tested five combinations of PRF and DC. For each pulsing protocol, we varied the pressure at the target between 0 and 2.2 MPa within a single experimental session with a randomized design (**Figure 2d**).

**Figure 2:**
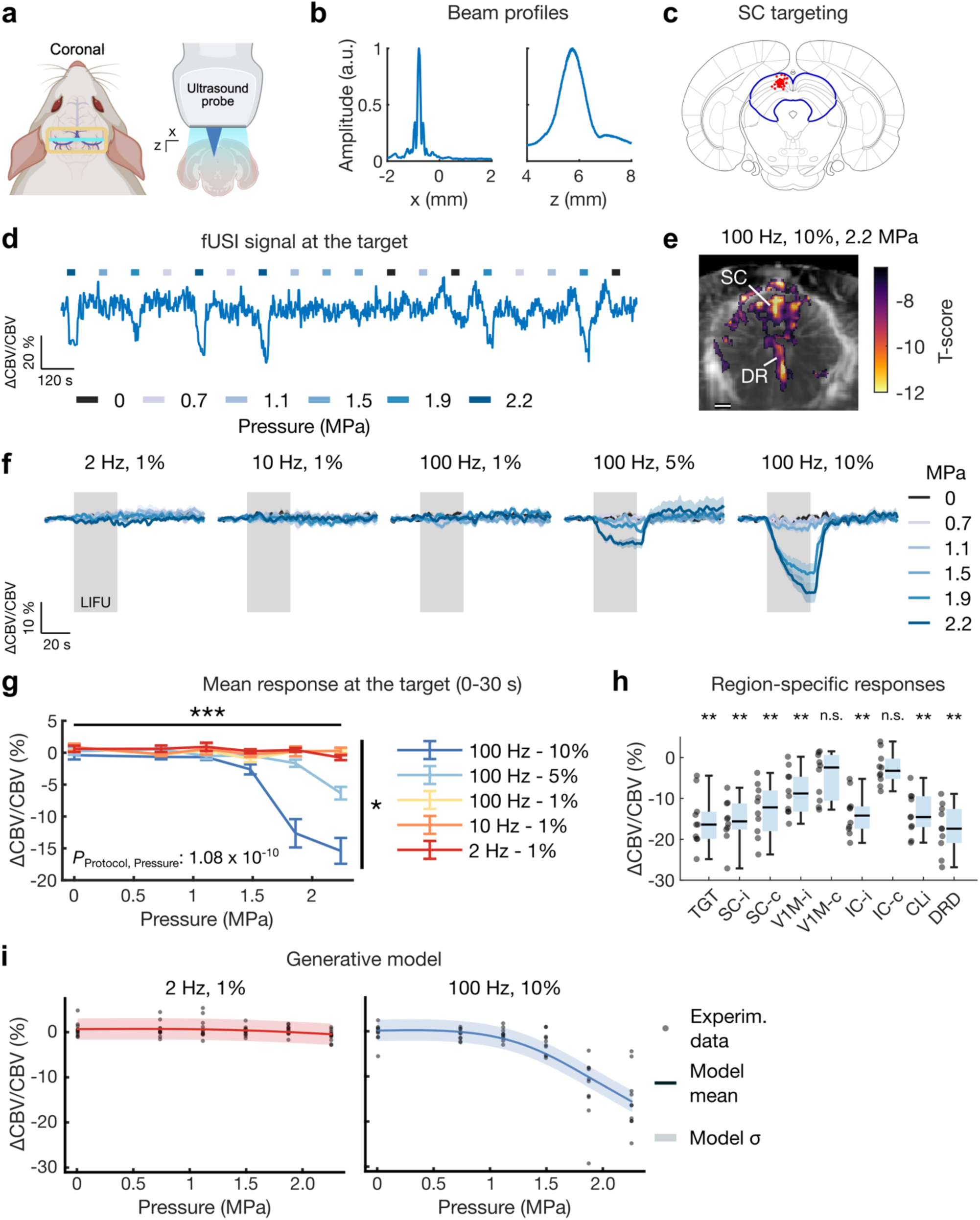
fUSI-LIFU parameter mapping. (**a**) Schematic of fUSI imaging in coronal view with LIFU stimulation of the rat superior colliculus (SC). (**b**) Beam profiles measured along the x and z directions at the SC target position. (**c**) LIFU target position in all the experimental sessions registered to the rat brain atlas (*N* = 48 sessions in 14 rats). The thick blue contour indicates the SC. (**d**) Representative fUSI time series at the SC target from an entire session of shuffled LIFU stimuli with different pressure values (3 repetitions per pressure). (**e**) LIFU-evoked fUSI responses for the configuration with pulse repetition frequency (PRF) of 100 Hz, duty cycle (DC) of 10%, and 2.2 MPa pressure overlaid on a representative power Doppler coronal image for orientation. Statistically significant T-scores (*P* < 10^-4^) were cluster-corrected to obtain *P*_FWER_ < 10^-4^. *N* = 10 rats. Scale bar: 1 mm. DR: dorsal raphe. (**f**) Stimulation-locked fUSI time series with different emitted pressures and pulsing protocols (defined as PRF-DC pairs). (**g**) Group-level analysis of the mean fUSI response in the 0-30 s interval during the LIFU stimulation. *P* = 0.01 (*) and *P* = 2.55 × 10^-11^ (***) in a mixed-effects model followed by ANOVA. *N* = 9-11 rats/group. (**h**) fUSI responses evaluated in brain regions adjacent to the target. *P* = < 0.01 (**) in a B-H–corrected Wilcoxon Rank Sum Test. TGT: target; V1M: monocular primary visual cortex; IC: inferior colliculus; Cli: caudal linear nucleus of the raphe; DRD: dorsal part of the dorsal raphe nucleus. (**i**) Generative model used for the in-silico BEACUN experiments in the SC based on the in-vivo parameter mapping. Only the cases for the (2 Hz, 1%) and (100 Hz, 10%) pulsing protocols are shown.

With the parameters tested, we only found putatively inhibitory responses, which appeared as a reduction in the ΔCVB/CBV signal (**Figure 2e**; *P*_FWER_ < 10^-4^). Specifically, we observed a large cluster of bilateral inhibition in the SC and a ventral cluster encompassing the raphe nuclei, which receive direct projections from the SC^34,35^. To quantify the response to each parameter configuration, we measured LIFU-evoked fUSI activity changes at the SC target (**Figure 2f-g**) and in adjacent brain regions (**Figure 2h**) time-locked to the stimulus onset. These responses varied with the pulsing protocol and dose-dependently increased as a function of the pressure. Specifically, we found significant effects for protocol (*F*_4,278_ = 3.29; *P* = 0.012), pressure (*F*_1,278_ = 48.35; *P* < 0.001), and interaction (*F*_4,278_ = 14.39; *P* < 0.001)

(**Figure 2g**).

### BEACUN hyperparameter tuning

Based on the results of the experimental fUSI-LIFU parameter mapping, we built a generative model that we used to carry out *in-silico* stimulation-response experiments (**Figure 2h**). These virtual experiments served as a platform to systematically determine the impact of relevant BO hyperparameters and to benchmark the performance of the BEACUN approach under different noise levels. To simulate noisy experimental conditions, we injected into the model white Gaussian noise with standard deviation equivalent to 1- or 2-fold the estimated noise standard deviation of the experimental ground-truth distribution (**Figure 2h**; **Supplementary Figure 1**). For each BO hyperparameter configuration, we carried out 50 independent optimization trials, each consisting of 100 evaluations, and benchmarked the BO performance against a random search.

The acquisition function is a critical component of the BO optimization loop, and different acquisition functions have been developed for noise-free and noisy environments^12^. We tested eight different acquisition functions to identify the one that would converge to the best solution in the least number of evaluations in our inhibitory fUSI-LIFU neuromodulation application. In the condition with 1-fold noise standard deviation, log-expected improvement^36^ (LogEI) and quasi-log noisy expected improvement^36^ (qLogNEI) achieved nearly equivalent results in terms of optimization regret and performed significantly better than the random search (*P* < 0.001) (**Figure 3a-c**). However, the LogEI acquisition function yielded a higher success rate than qLogNEI, with 96% of trials converging after 25 evaluations (**Figure 3d**).

**Figure 3:**
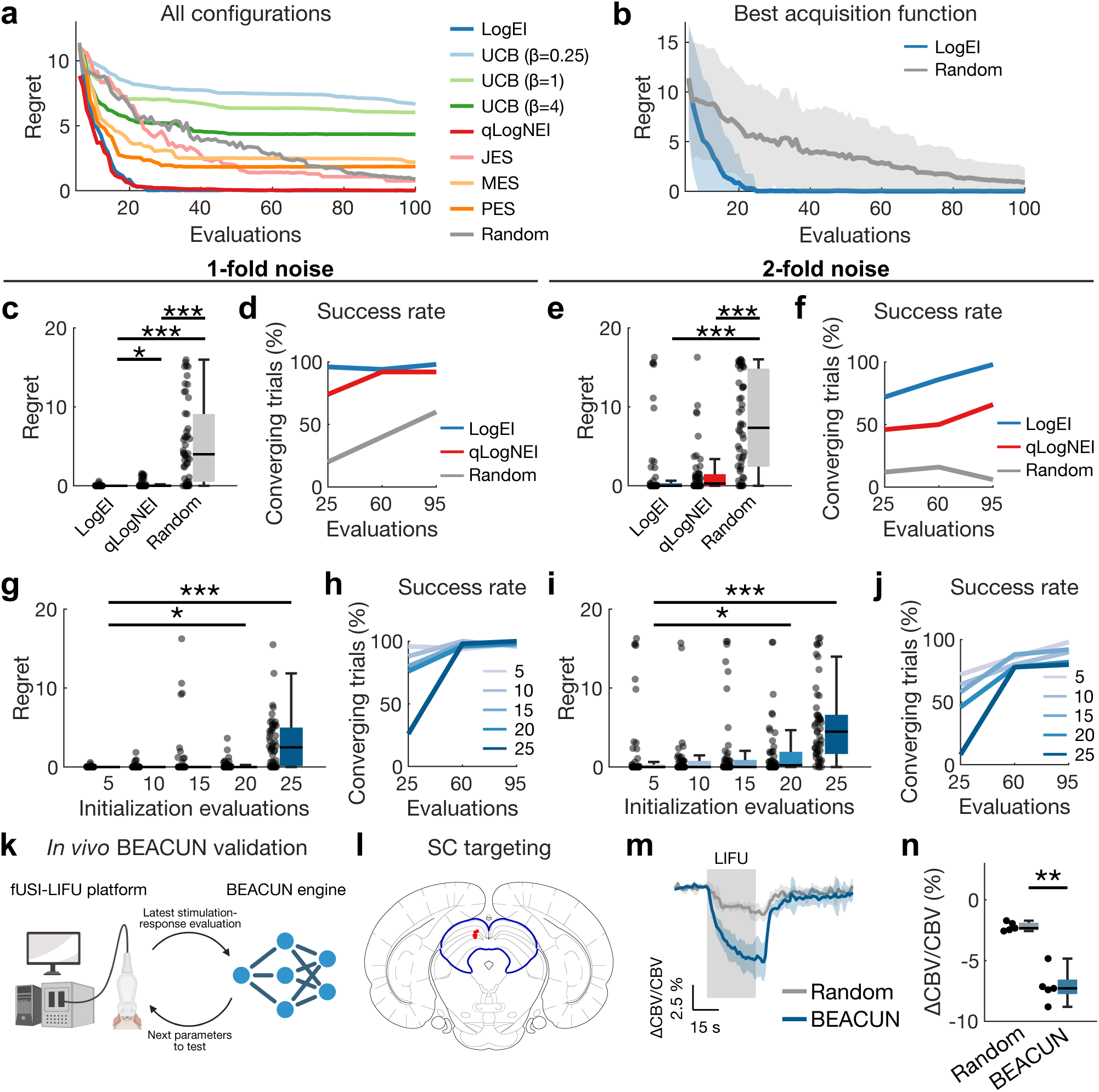
BEACUN hyperparameter tuning. (**a**) Optimization regret for all the BEACUN acquisition functions and random search with 1-fold noise. All the optimizations used 5 Sobol-distributed initialization points. The plots display the mean of 50 independent trials. (**b**) Regret time series for the optimization with the log-expected improvement (LogEI) acquisition function and random search. Solid lines display the mean and shaded areas show the standard deviation of 50 optimization trials. (**c**) Regret at evaluation 25 (5 initializations + 20 Bayesian optimization iterations) for the LogEI and quasi-log noisy expected improvement (qLogNEI) acquisition functions compared to the random search in the condition with 1-fold noise. LogEI vs. Random: *P* = 2.87 × 10^-13^ (***); qLogNEI vs. Random: *P* = 2.07 × 10^-10^ (***); LogEI vs. qLogNEI: *P* = 1.15 × 10^-2^ (*); B-H–corrected Mann-Whitney U-tests. (**d**) Success rate in the condition with 1-fold noise quantified as the number of trials (out of 50) converging with a regret lower than 1% of the ground-truth minimum. (**e**) Regret at evaluation 25 for the LogEI and quasi-log noisy expected improvement (qLogNEI) acquisition functions compared to the random search in the condition with 2-fold noise. LogEI vs. Random: *P* = 9.17 × 10^-9^ (***); qLogNEI vs. Random: *P* = 2.31 × 10^-8^ (***); LogEI vs. qLogNEI: *P* = 0.051 (n.s.); B-H–corrected Mann-Whitney U-tests. (**f**) Success rate in the condition with 2-fold noise. (**g**) Regret at evaluation 25 with varying Sobol-distributed initial points in the condition with 1-fold noise. We observed a significant effect for the number of initial points (*P* = 1.22 × 10^-36^ in a Kruskal-Wallis test). *P* = 0.03 (*) and *P* = 1.31 × 10^-11^ (***) in B-H–corrected Mann-Whitney U-tests. There was no significant difference between random and Sobol sampling (P = 0.20 in a Kruskal-Wallis test). (**h**) Success rate for the condition with 1-fold noise and varying Sobol-distributed initial points. (**i**) Regret at evaluation 25 with varying Sobol-distributed initial points in the condition with 2-fold noise. We observed a significant effect for the number of initial points (*P* = 7.41 × 10^-27^ in a Kruskal-Wallis test). *P* = 0.04 (*) and *P* = 1.96 × 10^-8^ (***) in B-H–corrected Mann-Whitney U-tests. There was no significant difference between random and Sobol sampling (*P* = 0.30 in a Kruskal-Wallis test). (**j**) Success rate for the condition with 2-fold noise and varying Sobol-distributed initial points. (**k**) Schematic representation of the BEACUN engine interfacing with the fUSI-LIFU platform in the in-vivo BEACUN optimization experiments. (**l**) LIFU target position registered to the rat brain atlas in the live BEACUN optimizations in vivo (*N* = 5 sessions in 5 rats). The thick contour indicates the SC region. (**m**) Stimulation-locked fUSI time series during the LIFU stimulation with the BEACUN and random searches. The plots show the parameter combinations that minimized the CBV response in each configuration after a total of 45 evaluations (5 initialization points + 40 search iterations). The pressure was limited to 1.6 MPa in these experiments. (**n**) Mean fUSI response in the 0-30 s interval for the random and BEACUN optimizations. *P* = 1.12 × 10^−3^ (**) in a paired two-tailed t-test. Hedge’s *g* effect size: -4.24; 95% confidence interval: (-6.5322, -1.8878).

Similarly, in the condition with 2-fold noise standard deviation, the LogEI and qLogNEI acquisition functions outperformed the random search (*P* < 0.001) (**Figure 3e**), but LogEI converged more quickly and at consistently higher rate (**Figure 3f**).

Next, we investigated the effect of the number and distribution of initialization evaluations ***X***_!"!#_. We repeated the optimizations with the LogEI acquisition function using *n*_!"!#_ between 5 and 25, and we compared random and Sobol sequence sampling. In both the conditions with 1- and 2-fold noise, we observed a significant effect for the number of initial points (*P* < 0.001) but found no differences between random and Sobol sampling (*P* > 0.20) (**Figure 3g, 3i, Supplementary Figure 5a-h**). We found converging evidence in the success rate analysis (**Figure 3h, 3j**). Finally, we evaluated the effect of different GP kernels, namely the radial basis function (RBF) and the 5/2 Matérn. We observed no difference in the 1-fold noise condition, whereas RBF performed better than Matérn in the 2-fold noise condition for the selected hyperparameters combination with *n*_!"!#_ = 5, Sobol sampling, and logEI acquisition function (*P* = 0.027) (**Supplementary Figure 6a-d**). Taken together, these results show that, given a fixed budget quantified in terms of stimulation-response evaluations, BEACUN with the LogEI acquisition function consistently outperformed random search and yielded better results when using the minimum number of initial evaluations. These results highlight the benefits of the smart parameter search guided by the acquisition function. These trends were especially prominent in the early phase of the search, i.e., within the first 60 evaluations, and are particularly relevant for real-world experimental conditions where scarce resources (e.g., limited budget, patient time, patient volumes) pose severe limitations to the number of feasible stimulation-response evaluations.

To further validate the findings of our *in-silico* hyperparameter tuning, we implemented a live BEACUN engine and carried out *in-vivo* experiments to benchmark its performance against a real-world random search (**Figure 3k**). We matched the experimental conditions to our ground-truth parameter mapping, and we directed LIFU stimuli to the SC target (**Figure 3l**). Importantly, in the live implementation we treated pressure, PRF, and DC as continuous parameters, expanding the search space and the complexity of the optimization relative to the *in-silico* experiments. After each stimulation, we quantified in real time the LIFU-evoked fUSI activity at the target time-locked to the stimulus onset (**Figure 3m**), and the mean ΔCVB/CBV value was used to update the GP model in the BEACUN engine. The parameters of the next LIFU stimulation were updated at runtime based on the next protocol candidate selected by the acquisition function. The live BEACUN implementation with 5 Sobol-distributed initialization points and LogEI acquisition function performed significantly better than the random search (*P* < 0.01; Hedge’s *g* effect size -4.24) and produced a ΔCVB/CBV signal reduction of -7.3% ± 1.19% (median ± IQR) (**Figure 3m-n**). These results provide a real-world validation of our *in silico* experimental setup and demonstrate that our BEACUN implementation generalized to an *in vivo* experimental setup.

### *In-silico* BEACUN validation with a behavioral LIFU stimulation model

We carried out additional *in-silico* experiments in our virtual environment to validate the performance of the BEACUN platform in a different LIFU stimulation application. We used a generative model based on real-world behavioral data recorded during LIFU stimulation of the centromedial thalamus (CMT) in rats^5^ (**Figure 4, Supplementary Figure 4a**). The search space included the acoustic pressure, the LIFU target (CMT or control region), and the stimulus duration (1 or 3 bursts). Unlike the SC objective function, which is largely monotonic with a boundary optimum (**Figure 2g-h**), the CMT landscape is non-monotonic with a local optimum (**Figure 4a**). This model allowed a meaningful characterization of the BEACUN optimization performance against a conventional grid search with increasing sampling density (**Figure 4b**). We introduced random displacements in the grid sampling in different trials to mimic realistic experimental conditions where the objective function is unknown and the sampled positions do not necessarily overlap with the ground-truth optimum (**Supplementary Figure 4b**).

**Figure 4:**
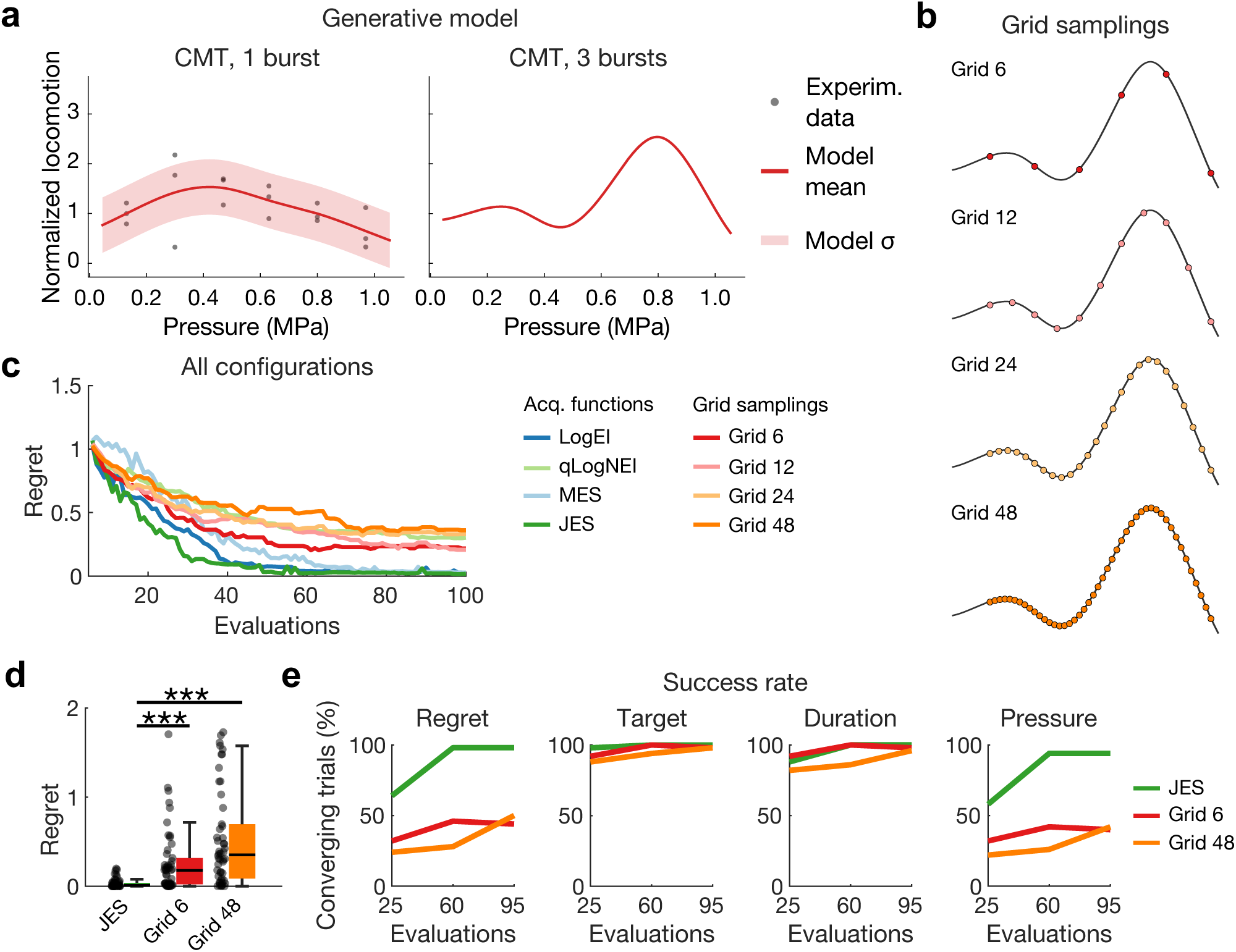
In-silico BEACUN optimization in a behavioral model of arousal. (**a**) Generative model used for the in-silico BEACUN experiments in the centromedial thalamus (CMT). Only the cases for the CMT target are shown. (**b**) Configurations used for the grid sampling approach with increasing number of samples. (**c**) Optimization regret for all the acquisition functions tested and comparison to the grid search with varying grid density. All the optimizations used 5 Sobol-distributed initialization points. The plots display the mean of 50 independent trials. (**d**) Regret at evaluation 50 (5 initializations + 45 Bayesian optimization iterations) for the JES and grid search with 6 and 48 samples. JES vs. Grid 6: *P* = 1.93 × 10^-6^ (***); JES vs. Grid 48: *P* = 8.55 × 10^-11^ (***); B-H–corrected Mann-Whitney U-tests. (**e**) Success rate quantified as 1) the number of trials (out of 50) converging with a regret lower than 5% of the ground-truth maximum; 2) the number of trials converging to the optimum target (CMT); 3) the number of trials converging to the optimum stimulation duration (3 bursts); 4) the number of trials converging within ± 5% of the optimum pressure.

In contrast to the SC environment, joint entropy search (JES) was the best-performing acquisition function in this environment (**Figure 4c**); furthermore, no significant differences among initialization strategies were observed here, unlike in the SC environment (**Supplementary Figure 5i-n**). These results reinforce the notion that optimization problems that appear similar and have comparable dimensionality and smoothness – e.g., SC and CMT objectives – may require fine-tuning of BO hyperparameters to achieve optimal performance. While JES performed significantly better than the grid search with all grid densities (*P* < 0.001) (**Figure 4d**), the convergence was slower than in the SC experiments (**Figure 4e**), possibly due to the higher complexity of the CMT optimization landscape (**Supplementary Video 1**). To further characterize the cause of this slower convergence, we calculated the success rate based on the optimum target, stimulus duration, and pressure. This analysis highlights that the pressure parameter is the main source of error in these optimization experiments. Interestingly, increasing the density of the grid sampling did not improve the performance of the grid search. A potential explanation for this behavior is that, while a finer sampling reduces the occurrence of systematic discretization errors, it exponentially increases the complexity of the optimization problem. Notably, these results show that BEACUN was able to effectively discriminate between the target of interest (CMT) and a control target located in a brain region decoupled from the neurobehavioral phenotype under investigation^5^. Using such control conditions is critical to ensure that the observed responses ensue from a direct LIFU-related mechanism and not from a nonspecific confounding effect^37–39^. Taken together, these results underscore the advantages of using a data-driven approach like BEACUN to effectively optimize LIFU neuromodulation protocols without relying on a fixed sampling of the parameter space determined a priori.

### Live BEACUN optimization of a LIFU protocol for suppression of event-related activations

To determine the performance of the BEACUN optimizer in the presence of event-related neural activities, we used a rat model of visual stimulation. In these experiments, we aimed to determine whether BEACUN could search for the optimal parameter configuration to inhibit the response to an external stimulus, akin to a symptom-specific neural biomarker. In addition, we implemented the probabilistic regret bound (PRB) stopping criterion to make the optimization fully automated^40^.

We performed binocular green light stimulation and directed concurrent blocks of LIFU stimuli to the thalamic dorsal lateral geniculate (DLG) nucleus (**Figure 5a-b**). At the baseline, visual stimulation produced significant activations in the DLG, primary visual cortex (V1), and postsubiculum (PoSub), consistent with previous reports^41^ (**Figure 5c**; *P*_FWER_ < 0.01). We carried out BEACUN optimization sessions in *N* = 5 rats, and in all sessions BEACUN was able to suppress visual-evoked responses measured by fUSI at the DLG target (**Figure 5d-f, Supplementary Figure 7, Supplementary Videos 2-3**). At the optimum, the PRF was 95.81 ± 9.38 Hz, the DC was 10%, and the pressure was 1 MPa (max pressure allowed in these experiments). The BEACUN-optimized LIFU stimulation produced a ΔCVB/CBV signal reduction of -5.7% ± 1% (median ± IQR) relative to the visual stimulation baseline (**Figure 5g**; *P* < 0.001; Hedge’s *g* effect size -12.68). Notably, the stopping criterion with a probability of success of 0.95 (P_95_) was met at evaluation 23 ± 3.67 (mean ± s.d.), which is a substantial budget reduction compared to the ground-truth parameter search (**Figure 5h**; *P* < 0.001). These results underscore the ability of our BEACUN approach to quickly and reliably identify effective LIFU stimulation parameters to inhibit event-related neural responses. In addition, our results demonstrate the ability of the BEACUN optimizer with the P_95_ probabilistic stopping criterion to carry out fully automated optimizations that stop when further stimulation-response evaluations are deemed of marginal benefit. Altogether, our findings demonstrate the potential of the BEACUN optimizer to effectively identify LIFU protocols in real-world experimental settings in a variety of brain states.

**Figure 5:**
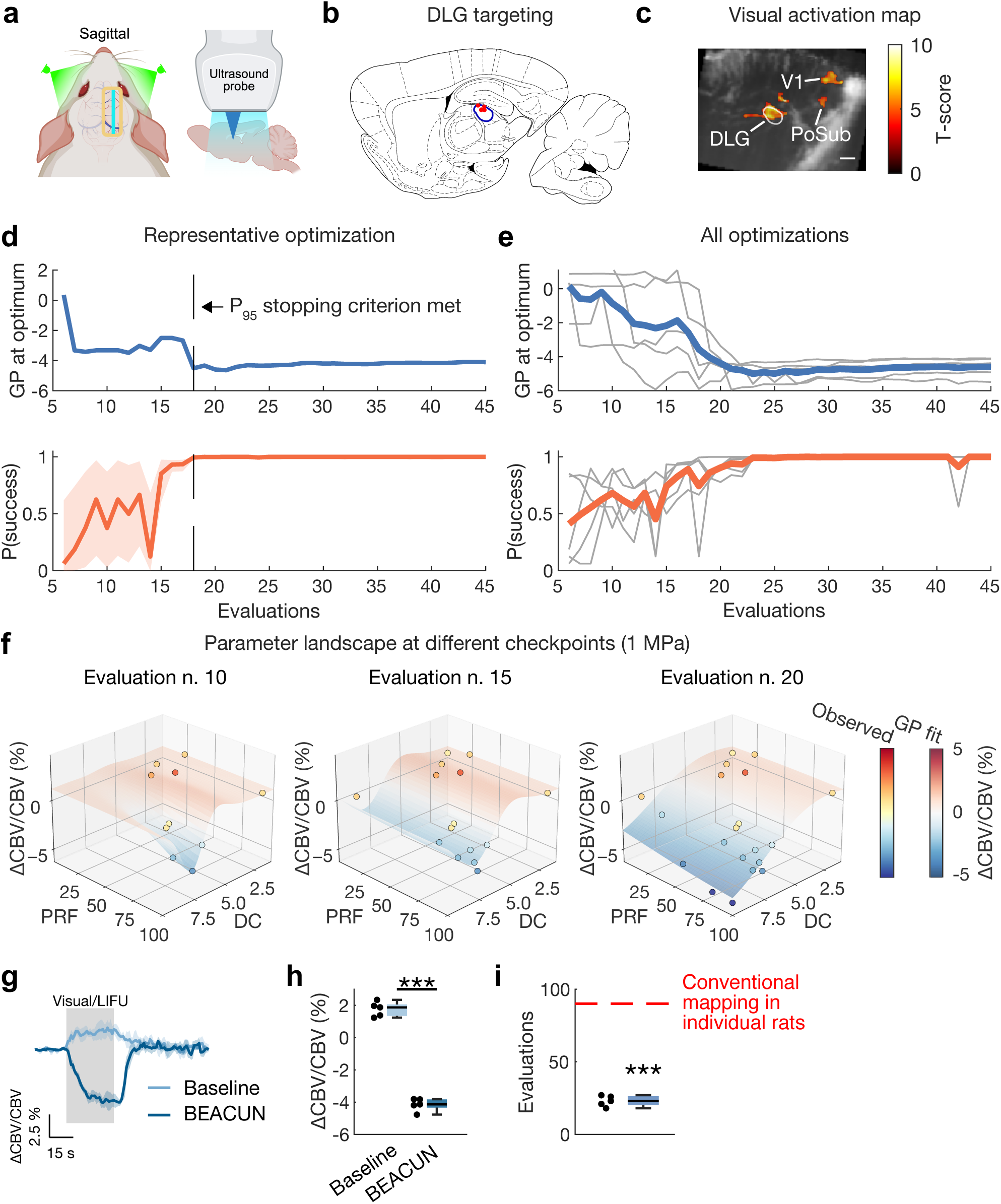
Live BEACUN optimization of inhibitory LIFU for suppression of visual-evoked activations in the dorsal lateral geniculate nucleus. (**a**) Schematic of fUSI imaging in sagittal view with concurrent visual stimulation and LIFU targeting of the thalamic dorsal lateral geniculate (DLG) nucleus. (**b**) Target position of the LIFU stimulation registered to the rat brain atlas (*N* = 5 rats). The thick blue contour indicates the DLG region. (**c**) Visual-evoked fUSI responses overlaid on a representative power Doppler sagittal image for orientation. *P*_FWER_ < 10^-2^. Scale bar: 1 mm. V1: primary visual cortex. PoSub: postsubiculum. (**d**) Representative time series showing the temporal evolution of the Gaussian process (GP) optimum and probability of success. The orange shaded area represents the 95% confidence interval of the success probability. In this experiment, the P_95_ stopping criterion was met after 18 evaluations (5 initialization points + 13 BO iterations). (**e**) Group-level time series of GP optimum and probability of success. Gray lines display the individual sessions. Thick colored lines display the group-level means. (**f**) Parameter landscape showing the evolution of the BEACUN search at relevant checkpoints. (**g**) Stimulation-locked fUSI time series during the visual/LIFU stimulation for the baseline and optimum conditions. The pressure was limited to 1 MPa in these experiments. (**h**) Mean fUSI response in the 0-30 s interval evaluated at the baseline and optimum. *P* = 7.16 × 10^−5^ (***) in a paired two-tailed t-test. Hedge’s *g* effect size: -12.68; 95% confidence interval: (-18.86, -6.5). (**i**) Number of BEACUN evaluations (including 5 initialization points) to reach the P_95_ stopping criterion compared to the number of trials (90; red dashed line) performed in individual rats in the grid-search parameter mapping. *P* = 2.16 × 10^−6^ (***) in a one-sample two-tailed t-test.

## Discussion

LIFU neuromodulation is an emerging modality for the noninvasive treatment of neuropsychiatric conditions. Preliminary evidence of LIFU’s potential has been reported in applications spanning modulation of emotional processing for mood disorders^3^, reduction of beta-band power in Parkinson’s disease^42^, modulation of sleep patterns^43^, and reduction of depression symptoms^1^. Despite these advances, the LIFU parameter space remains poorly characterized, and additional challenges related to the high variability in the treatment response^7^, precise target identification in each subject^2^, and lacking confirmation of target engagement continue to slow clinical adoption of LIFU stimulation.

Closed-loop parameter mapping is an impactful approach that aims to identify optimal settings for the patient-specific neurobiology and symptom profile by testing different stimulation configurations while monitoring the ensuing neural responses^9^. Variations of this approach have been successfully deployed in deep brain stimulation in major depressive disorder, where precise circuit-targeted protocols have been developed through expansive individualized stimulation-response mapping^10,44^. Similarly, in transcranial magnetic stimulation, the target location can be refined based on patient-specific resting-state functional magnetic resonance imaging (fMRI) connectivity^11^. Nevertheless, optimizing stimulation parameters and brain targets remains challenging and costly. Furthermore, time-dependent factors like disease progression and neuroplastic adaptations to the stimulation may decrease the therapeutic efficacy over time, making repeated optimizations necessary^45,46^.

To enable closed-loop parameter mapping in LIFU neuromodulation, we developed BEACUN, a method for data-driven, adaptive, and automated parameter exploration. BEACUN significantly reduces the cost of the optimization routine while converging to better and more robust stimulation protocols. We carried out extensive validations *in silico* and *in vivo* to tune relevant hyperparameters and benchmarked the optimization performance against conventional parameter exploration methods. Our results demonstrate that BEACUN provides a framework for effective stimulation-response mapping that could pave the way for personalized LIFU protocol optimization in clinical populations. While in our live BEACUN experiments we primarily focused on temporal pulsing parameters like the PRF, DC, and stimulus duration, other parameters like the brain target could be included in the optimization, as demonstrated by our CMT experiments. In the future, this open-target approach could be used for unbiased, brain-wide stimulation-response mapping integrating active control targets^1,5^.

Combining LIFU with concurrent functional neuroimaging provides a unique opportunity to confirm target engagement by monitoring real-time stimulation-evoked responses in a wide field of view. This has been demonstrated using Ca^2+^ imaging^6,47^ and photoacoustic tomography^48^ in rodents and fMRI in humans^2^. We based our live BEACUN experiments on fUSI measures to capitalize on the high sensitivity of this modality and to leverage its ease of use in rodent models^30,49^. Our fUSI-LIFU implementation realizes a robust and flexible platform that is easily reproducible, as it uses only off-the-shelf ultrasound components, and could promote the development and validation of other novel neuroimaging-based methods.

Another important component of our BEACUN design is the rule used to terminate the optimization. Prior optimization studies based on Bayesian approaches have generally relied on simple heuristics to halt the search, e.g., stopping after a fixed budget is exhausted or after the recommended optimum has not changed for a predetermined number of consecutive iterations^20,26,28^. In contrast, we used the PRB criterion^40^, a principled approach that bases the stopping decision directly on the GP posterior.

There are several limitations to this study. First, the high frequency used in the fUSI-LIFU experiments differs from transcranial LIFU applications in humans, which typically use sub-MHz LIFU stimuli. However, we demonstrated that the BEACUN optimization performance generalized well across experimental settings independent of the specific setup and stimulation-response function. This was the case under both *in silico* and *in vivo* conditions. In addition, the high frequency produces a narrow beam profile (0.1-0.2 mm) that is better suited for experimentation in the rodent brain. Our fUSI-LIFU findings are also directly relevant to intracranial LIFU applications where higher stimulation frequencies can be used for increased spatial specificity. Another limitation is that in the fUSI-LIFU experiments we quantified cerebrovascular neural correlates, which are an indirect measure of neural activity. While previous studies have confirmed the neural basis of the fUSI signal^30,50^, we cannot rule out a direct vascular effect of the LIFU stimuli.

In summary, our results establish BEACUN as a platform for efficient, data-driven optimization of LIFU neuromodulation parameters. Our live BEACUN implementation converged to the optimal LIFU stimulation protocol after only 23 ± 3.67 stimulation-response evaluations, and it performed significantly better than conventional parameter mapping methods in all the experimental settings tested. Importantly, BEACUN addresses two key limitations that are currently slowing the development of clinically relevant LIFU neuromodulation protocols, i.e., the poorly characterized parameter space and the high inter- and intra-subject response variability. In doing so, BEACUN could pave the way for personalized, subject-specific LIFU optimization.

## Methods

### Acoustic field characterization

To characterize the acoustic field of the LIFU emissions, we performed hydrophone measurements using an HGL-0085 hydrophone (Onda Corporation, Sunnyvale, CA, USA) immersed in degassed water. The field was scanned using a 3-axis positioning system (AIMS III, Onda Corporation), and the precise hydrophone position was determined based on the ultrasound time of arrival. Beam profiles were scanned with a resolution of 10 μm. At the SC target, the measured focal full-width at half maximum (FWHM) was 0.13 mm laterally, 0.41 mm in the elevation direction, and 1.18 mm along the depth. At the DLG target, the FWHM was 0.23 mm laterally, 0.69 mm in the elevation direction, and 2.58 mm along the depth.

For the voltage-pressure calibrations at the SC and DLG target positions, we measured the peak-negative pressure at the spatial peak position as a function of the Verasonics transmit power controller (TPC) voltage. The TPC voltage was varied between 1.6 and 8 V in steps of 0.2 V. The pulse length was adjusted to ensure that a steady state was reached while avoiding reflections between the transducer and hydrophone. Each measurement was repeated three times to filter out experimental noise. We derated the measured pressure by 0.3 dB cm^-1^ MHz^-1^. In the fUSI-LIFU parameter mapping, the acoustic pressure was limited to 2.2 MPa (MI = 0.56). In the live BEACUN experiments, the acoustic pressure was limited to 1.6 MPa (SC; MI = 0.41) or 1 MPa (DLG; MI = 0.26).

### Animals

All animal procedures were approved by the Institutional Animal Care and Use Committee at the University of California San Francisco. We used male and female Long Evans rats that were 8-10 weeks old at the time of the surgery. Rats had *ad libitum* access to water and food for the entire duration of the experiments and were housed in a temperature-controlled vivarium on a 12-h light-dark cycle (lights on at 7 AM; lights off at 7 PM). Following the surgical procedures, we maintained postoperative analgesia with buprenorphine (0.1 mg/kg; subcutaneously). Rats were singly housed and were allowed to recover for at least 7 days before any further experimentation.

### Surgical preparation

Rats received a surgical craniotomy and chronic prosthesis implantation as described previously^30^. Briefly, the animals were kept under isoflurane anesthesia (3.5% induction; 1% maintenance) in a digital stereotaxic frame for head fixation and orientation (Stoelting Co.), and body temperature was maintained at 37 °C using a warming pad (RightTemp Jr.; Kent Scientific). The incision site was shaved and disinfected by applying povidone-iodine and 75% EtOH. For imaging in coronal view, the cranial window extended from bregma -3 mm to bregma -9 mm AP, with an ML span of ±6 mm. For imaging in sagittal view, the cranial window extended from bregma to bregma -9 mm AP and from bregma +1 mm to bregma +5 mm ML. The bone was pretreated using a bonding agent (iBOND Total Etch; Kulzer). Bone flaps were cut using a high-speed drill with a 0.7-mm drill bit (Fine Science Tools) and were gently removed avoiding any damage to the dura mater. The craniotomy was then covered using a 125-μm polymethylpentene film sealed with dental cement (Tetric EvoFlow; Ivoclar Vivadent). The space between the dura and the film was filled with 0.9% sterile saline.

### fUSI-LIFU parameter mapping

#### fUSI-LIFU acquisition

We implemented the fUSI-LIFU sequence on a Vantage NXT 256-channel research scanner (Verasonics Inc., Kirkland, WA, USA) connected to a linear array transducer (Vermon, France; 15-MHz center frequency; 128 elements; 0.1-mm pitch). We used centrifuged ultrasound gel for acoustic coupling. The fUSI sequence consisted of nine tilted plane waves with uniformly distributed angles in the -8° to 8° range emitted with a pulse repetition frequency of 24 kHz. The frames were beamformed in a regular grid of pixels with in-plane resolution of 0.1 × 0.1 mm^2^. Beamforming was performed in real-time in an NVIDIA RTX A6000 GPU.

We used 200 compounded frames per power Doppler image acquired at a rate of 1 kHz. To eliminate the clutter signal originating from the tissue, we used a 5^th^-order temporal high-pass Butterworth filter with a cutoff frequency of 40 Hz and a singular value decomposition filter that eliminates the first singular value^17,30^. The power Doppler intensity was computed at each pixel, yielding a final frame rate of approximately 1 frame/s.

#### fUSI-LIFU pulsing sequence

We used the same ultrasound probe to acquire fUSI data and to transmit interleaved LIFU emissions. LIFU transmit events consisted of focused beams emitted by a 6-mm sub-aperture. The target coordinates were defined based on the rat brain atlas^51^. To increase the volume of brain tissue engaged by the LIFU stimuli, we used two foci laterally displaced around the nominal target by 500 μm. Transmit events for the two foci were interleaved to ensure that the same acoustic dose was delivered at each location. To prevent transducer overheating, the duration of individual LIFU pulses was limited to 5 ms based on thermal measurements performed using an infrared camera.

To map the parameter space of the fUSI-LIFU stimulation in the SC target, we used repeated stimulation sessions during which we tested the following protocols defined as combinations of PRF and DC: 1) 100 Hz, 10%; 2) 100 Hz, 5%; 3) 100 Hz, 1%; 4) 10 Hz, 1%; 5) 2 Hz, 1%. In each session, we fixed the PRF and DC and varied the pressure in six steps between 0 and 2.2 MPa. For each pressure, we performed three trials. The order of the trials with the different LIFU protocols was shuffled within individual sessions, and the order of the protocols was shuffled at the animal level to control for order and carry-over effects.

Each stimulation trial lasted 110 s and consisted of three phases: a 20-s baseline, a 30-s period of interleaved fUSI and LIFU stimuli, and a 60-s post-stimulation period. During the 30-s stimulation period, a 500-ms LIFU pulse train was delivered (alternatively between the two foci), followed by a 750-ms delay before the next fUSI acquisition to minimize interference between the two modalities. The frame rate during the 30-s stimulation period was approximately 0.5 frame/s.

#### fUSI data processing

We registered the fUSI frames from each session through rigid transformations to a high-resolution intra-session template calculated as the median of 50 fUSI images. Motion artifacts were filtered in each data set following previously described protocols^30^. Each fUSI acquisition was then manually aligned to the brain atlas^51^ using anatomical landmarks and was temporally interpolated to obtain a uniform frame rate (1 Hz).

We segmented the target as a rectangular region of interest (ROI) of 0.63 (lateral) × 1.18 (axial) mm^2^ in the SC experiments and 0.73 (lateral) × 1.5 (axial) mm^2^ in the DLG experiments. The ROI lateral dimensions were determined based on the lateral FWHM and lateral focus displacement; the axial ROI dimensions were determined based on the axial FWHM in the SC experiments and the axial span of the target region in the DLG experiments. We time-locked the CBV signals from each trial to the LIFU stimulation onset, then we calculated the ΔCBV/CBV signals from each pre-LIFU 20-sec baseline. The signals were temporally smoothed by a moving average filter (3 time points).

For the statistical analyses comparing the effects of the different pulsing protocols and emitted pressures, we quantified the mean response in each animal and each session (average of three repetitions) during the 0-30 s time interval. Then, a mixed-effects model was fitted to the mean responses using *protocol* and *pressure* as fixed effects and *subject* and *session* as random effects using the MATLAB fitlme.m function and the following model design:

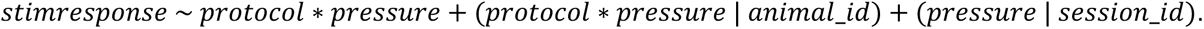

The mixed-effects model was followed by ANOVA to evaluate the significance of the fixed-effects terms and interactions.

#### fUSI-LIFU functional maps

To generate the functional maps, we first quantified the pixel-wise mean from the stimulation-locked ΔCBV/CBV signals in the 0-30 s time interval (average of three repetitions per session). Then, we applied spatial smoothing using a Gaussian filter with a FWHM of 0.3 × 0.3 mm^2^, and we performed pixel-level inference by calculating the group-level t-scores. We masked the pixel values for statistical significance at *P* < 10^-4^ (two-tailed) and applied a cluster correction threshold of 13 pixels (*P*_FWER_ < 10^-4^) calculated using the 3dClustSim.m program of the AFNI library^52^.

### BEACUN implementation

#### Gaussian process surrogate model

The surrogate model was a Gaussian process (GP) implemented via BoTorch. For the live experiments, a SingleTaskGP was used since the search space was treated as continuous. For the offline simulations, a MixedSingleTaskGP was used to handle the mixed categorical-continuous input space. In both cases, the continuous dimension was normalized to the [0, 1] range and observed responses were standardized to zero mean and unit variance. We evaluated two kernels for the continuous component of the GP: a radial basis function (RBF) kernel and a 5/2 Matérn kernel. The RBF kernel assumes infinitely differentiable sample paths, whereas the Matérn kernel provides a less smooth prior over functions. Both used automatic relevance determination (ARD) lengthscales. For the SingleTaskGP, this kernel acted directly on the continuous inputs. For the MixedSingleTaskGP, it was combined with a categorical kernel (also using ARD) and was defined as:

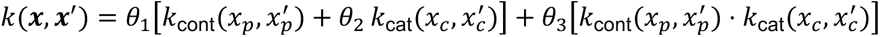

where *x_p_* and *x_c_* are the continuous (pressure) and categorical (parameter condition) components, *θ*_1_, *θ*_2_, *θ*_)_ are learned scale parameters, *k*_cont_ is a continuous kernel over the pressure dimension, and *k*_cat_ is a categorical kernel. The additive component captures marginal effects of pressure and condition independently, while the interaction component captures condition-dependent pressure responses. The lengthscale prior was set to the BoTorch default (LogNormal), which is scaled to dimensionality^53^. The noise variance prior was set to the legacy BoTorch Gamma (1.1, 0.05), which is nearly uninformative and allows the noise level to be determined primarily by the data. GP hyperparameters were optimized at each iteration by maximizing the exact marginal log-likelihood via L-BFGS-B with multiple random restarts.

#### Acquisition functions

We evaluated 8 acquisition functions that we considered as hyperparameters:

##### Log-expected improvement (LogEI)

The expected improvement (EI) quantifies the expected amount by which a new observation will improve upon the current best. LogEI computes EI in log-space, which provides better numerical stability when the improvement probability is very small^36^.

##### Upper confidence bound (UCB)

UCB selects the point that maximizes 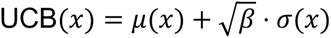, where *μ*(*x*) and *σ*(*x*) are the GP posterior mean and standard deviation (for minimization problems, the corresponding lower-confidence-bound form was used). The parameter *β* controls the exploration–exploitation trade-off: larger *β* favors querying uncertain regions (exploration), while smaller *β* favors querying near the current best estimate (exploitation). Three settings were evaluated: *β* = 0.25 (exploitation-focused), *β* = 1.0 (balanced), and *β* = 4.0 (exploration-focused).

##### Monte Carlo noisy expected improvement (qLogNEI)

Standard EI assumes the current best observation is known exactly, but under noisy observations this information is not available. qLogNEI addresses this by computing a Monte Carlo estimate of the expected improvement, marginalizing over the uncertainty in the current best^36^.

##### Max-value entropy search (MES)

MES is an information-theoretic acquisition function that selects the point maximizing the mutual information between the next observation and the unknown maximum value *f*^∗^ of the objective^54^. The mutual information is approximated via a Gumbel-based lower bound.

##### Predictive entropy search (PES)

PES maximizes the mutual information between the next observation and the optimal input location *x*^∗^. The acquisition value is computed via expectation propagation (EP), which approximates the intractable posterior over the optimum location^55^.

##### Joint entropy search (JES)

JES extends entropy-based approaches by maximizing the mutual information between the next observation and the joint distribution of the optimal input-output pair (*x*^∗^, *f*^∗^). Optimal input-output samples were obtained from the GP posterior, and entropy was estimated using a lower-bound estimator^56^.

#### Live BEACUN optimization

For the live BEACUN experiments, we implemented a real-time fUSI processing pipeline and quantified the ΔCBV/CBV signal after each LIFU stimulation trial in an ROI centered at the LIFU target location (SC or DLG). The mean ΔCBV/CBV during the 30-s stimulation period was fed into the BEACUN engine to update the GP regression and obtain the parameters for the next LIFU stimulation. The LIFU parameters were then updated at runtime in the Verasonics fUSI-LIFU platform before the next stimulation was delivered.

#### Stopping Criterion

We implemented the probabilistic regret bound (PRB) stopping criterion^40^. PRB stops the optimization when a candidate point ***X*** is deemed (*ε*, *δ*)-optimal, i.e., the probability that its regret is at most *ε* exceeds 1 − *δ* under the GP posterior. At each iteration *t*, the posterior GP model was used to draw approximate function samples from which the model-based regret 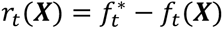 was estimated yielding Monte Carlo estimates Ψ_*t*_(*x*) = *P*(*r*_*t*_(***X***) ≤ *ε*). To guard against premature stopping due to Monte Carlo estimation error, Clopper-Pearson confidence intervals were used to ensure that Ψ_*t*_(***X***) is estimated with sufficient precision. The total error tolerance *δ* is partitioned between a model error component *δ*_mod_ and an estimation error component *δ*_est_, ensuring that any point passing the stopping criterion is *ε*-optimal with probability at least 1 − *δ*.

In our live experiments, we fixed the total error tolerance at *δ* = 0.05, split equally between *δ*_mod_ and *δ*_est_. The estimation budget *δ*_est_ was divided uniformly across iterations via a constant schedule, allocating each iteration an equal share. The regret tolerance *ε* was set at 0.5% ΔCVB/CBV signal change.

#### Benchmarks

We used two benchmark strategies to provide reference performance levels against which different BO arms were compared. In the random search, a query point was drawn from a uniform random distribution at each optimization iteration. A GP was fit at each step to estimate the current optimum via the posterior mean but was not used to guide point selection. In the grid search, the pressure domain was discretized into an equally spaced grid within each condition, with a random proportional offset drawn once for each trial. At each optimization iteration, a random pair was queried. The optimum was estimated as the grid coordinate with the best average noisy response. We evaluated three grid densities: 6, 12, 24, and 48 points per condition.

### Live BEACUN optimization in the DLG target

#### Visual stimulation

We performed binocular green light stimulation as reported previously^17^. Briefly, anesthetized rats were habituated in a dark chamber for at least 30 min prior to visual stimulation. Bilateral stimulation blocks were delivered using green-light LEDs controlled by a microcontroller board (Arduino Uno) interfaced with the Verasonics scanner for precise synchronization with the imaging sequence. For each stimulation block, the LEDs were flashed for 30 s at a frequency of 3 Hz with a 60-s pause. We recorded fUSI data continuously in sagittal view (bregma +3.4 ML). Five initial blocks were used image the baseline response to the visual stimulation.

#### Visual activation maps

To generate the visual activation maps, we performed pixel-level inference using a general linear model (GLM) implemented in MATLAB based on the glmfit.m function. We calculated the ΔCBV/CBV signals from an initial 20-sec baseline. The signals were detrended using a 4^th^-order polynomial fit and were temporally smoothed by a moving average pixel-wise filter (3 time points). We applied spatial smoothing using a Gaussian filter with a FWHM of 0.3 × 0.3 mm^2^. We generated the activation model regressor for the GLM by convolving the stimulation blocks (visual) with a single-gamma hemodynamic response function (HRF)^57^. We used an HRF time constant of 0.7, a time delay of 1 s, and a phase delay of 3 s (Ref. ^58^). To compute the group-level functional maps, we performed 2^nd^-level inference by calculating the t scores from the GLM Δ values. We masked the pixel values for statistical significance at *P* < 10^-2^ (two-tailed) and applied a cluster correction threshold of 25 pixels (*P*_FWER_ < 10^-2^) calculated using the 3dClustSim.m program of the AFNI library^52^.

### Software

All BEACUN experiments were carried out in Python. GP models and acquisition functions were built using BoTorch^59^ and GPyTorch^60^. The probabilistic regret bound stopping criterion, originally implemented in Trieste^40,61^, was translated to BoTorch.

## Acknowledgements

We would like to thank Prof. Andrew Krystal, Prof. Leo Sugrue, Prof. Khaled Moussawi, and all members of the UCSF Focused Ultrasound in Neuroscience Program for the support and for engaging discussions on LIFU neuromodulation and LIFU parameter mapping. We would like to express our appreciation to Prof. Joshua Berke for the continued support to our lab. We thank all present and past members of the Di Ianni Lab for thoughtful discussions and feedback on this work.

## Author Contributions

Study conception, design, and supervision: T.D.I. Funding acquisition: T.D.I. Data collection: V.G. and A.B. Interpretation of results: A.B., V.G., and T.D.I. Paper preparation: T.D.I. and A.B. All authors reviewed the results and approved the final version of the manuscript.

## Data and materials availability

Data and codes will be released on Figshare and GitHub at the time of publication.

## Supplementary Figures

**Supplementary Figure 1.**
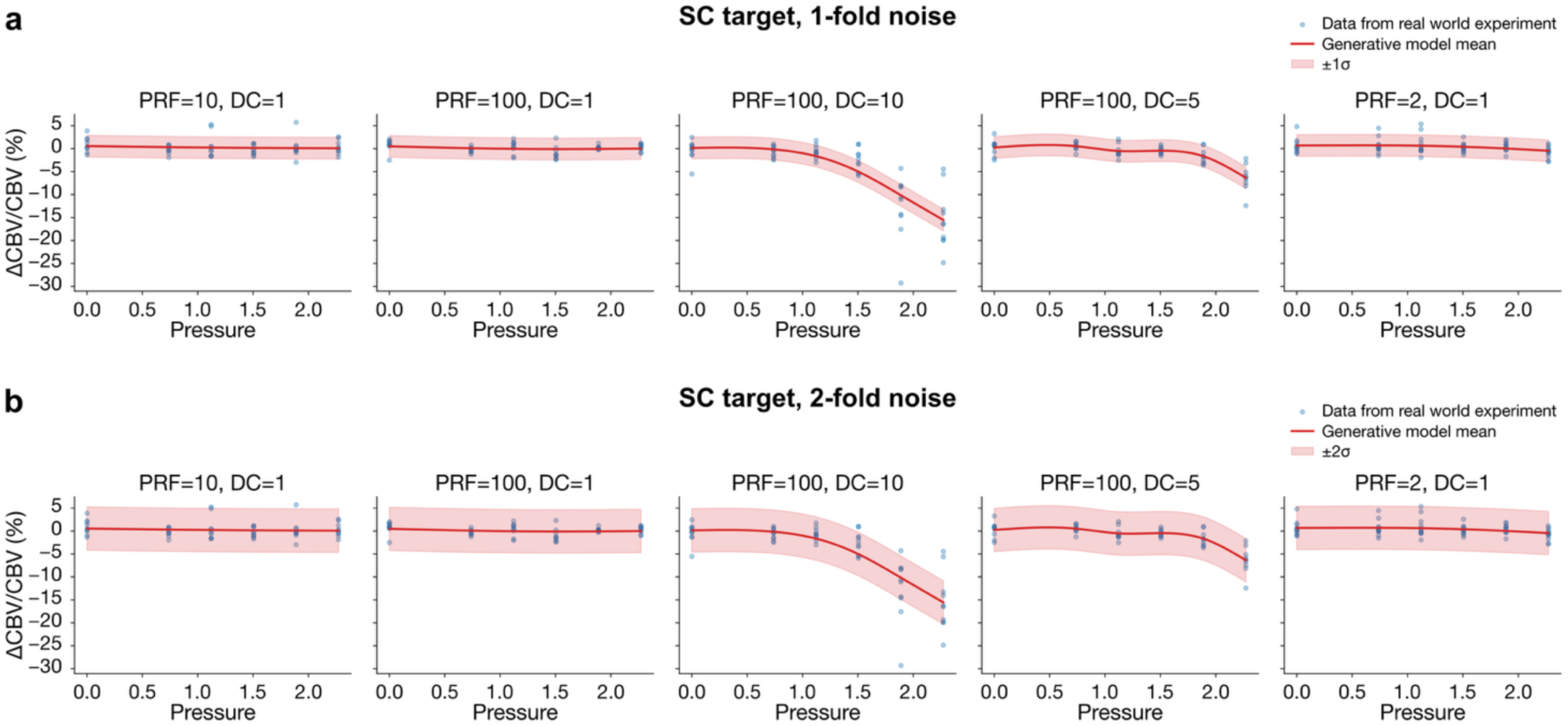
Generative model for the *in-silico* experiments in the superior colliculus (SC) target with 1-(**a**) and 2-fold (**b**) noise standard deviation.

**Supplementary Figure 2.**
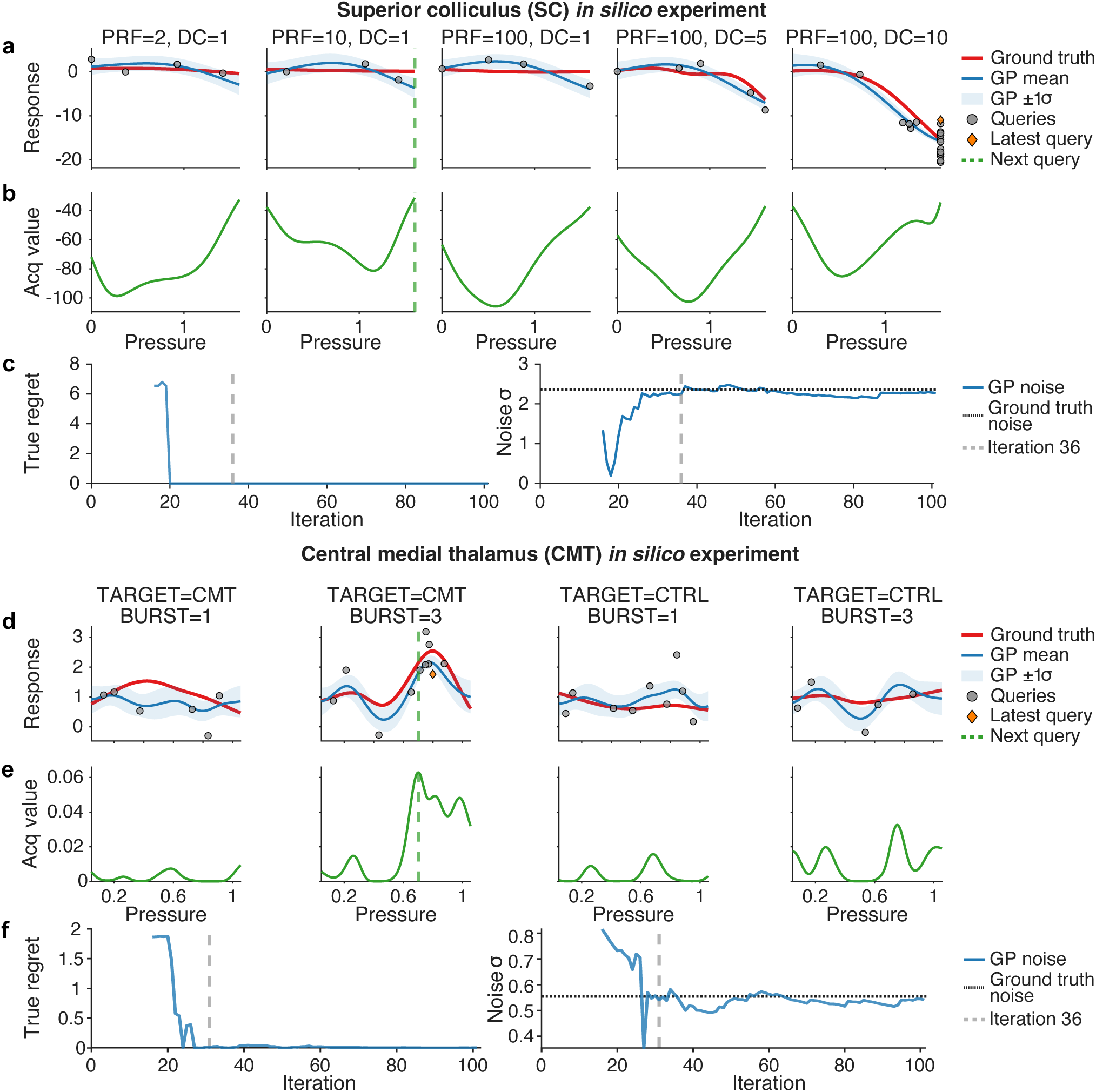
Representative iteration from one trial of the SC (**a-c**) and CMT (**d-f**) *in silico* experiments. Panels (**a,d**) show the ground-truth objective function (red), together with the GP posterior mean and ±1 SD uncertainty band (blue) fitted to the queried points up to the iteration shown. Queried points are shown in grey, and the latest query is shown in orange. Panels (**b,e**) show the acquisition function computed from the corresponding GP surrogate; the next query (dashed green line) is obtained by maximizing the acquisition function. Panels (**c,f**) show, on the left, the regret progression over the full trial, with the grey dashed line marking the displayed iteration. On the right, panels show the GP-estimated noise σ, computed as the square root of the fitted likelihood noise variance; the dashed black line indicates the ground-truth noise level used for the experiment.

**Supplementary Figure 3.**
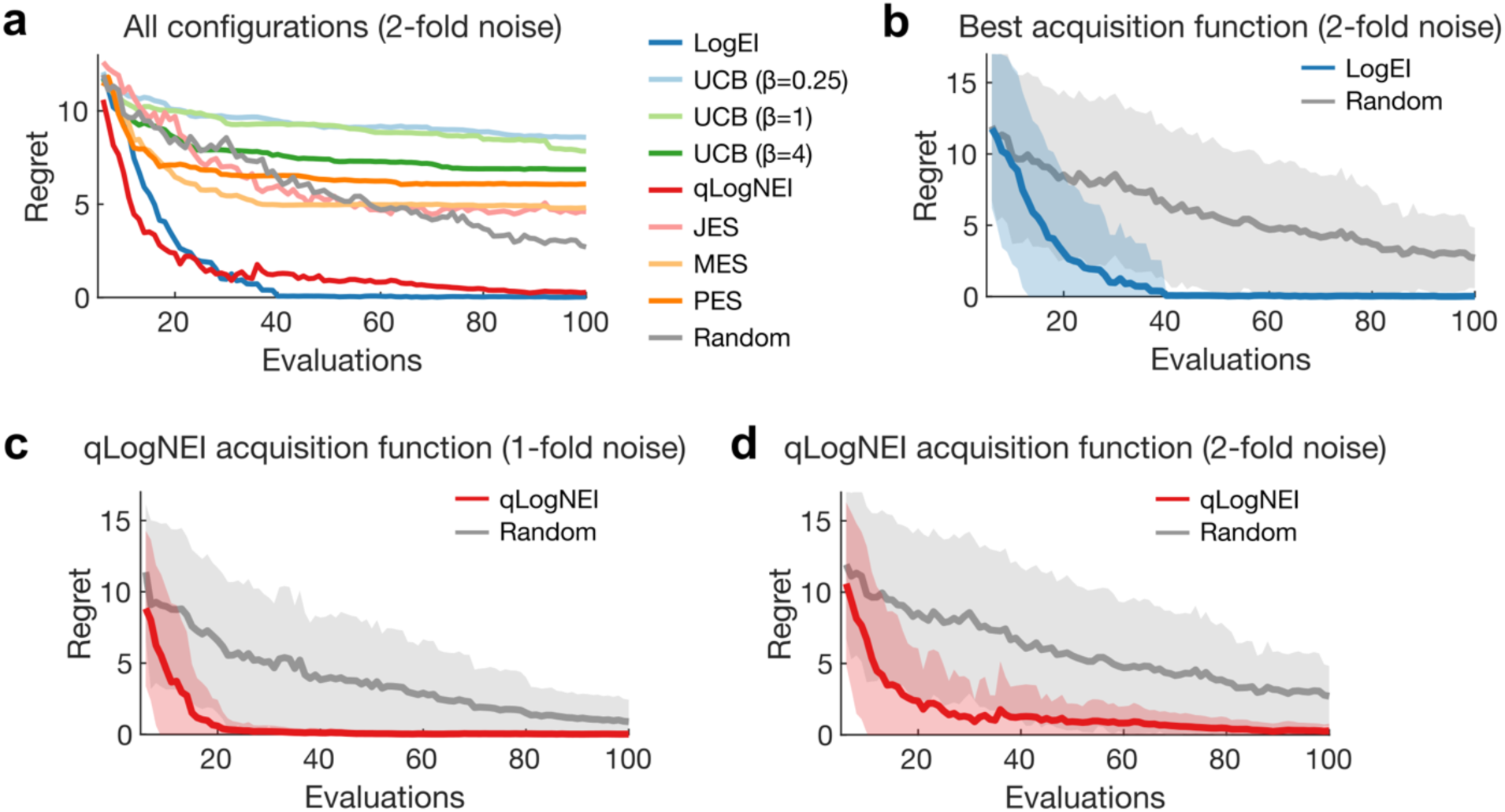
(**a**) Optimization regret for all the acquisition functions tested and random search with 2-fold noise. All the optimizations used 5 Sobol-distributed initialization points. The plots display the mean of 50 independent trials. (**b**) Regret time series for the optimization trials with the log-expected improvement (LogEI) acquisition function and random search with 2-fold noise. Solid lines display the mean and shaded areas show the standard deviation of 50 optimization trials. (**c, d**) Regret time series for the optimization trials with the quasi-log noisy expected improvement (qLogNEI) acquisition function and random search with 1-(**c**) and 2-fold (**d**) noise. Solid lines display the mean and shaded areas show the standard deviation of 50 optimization trials.

**Supplementary Figure 4.**
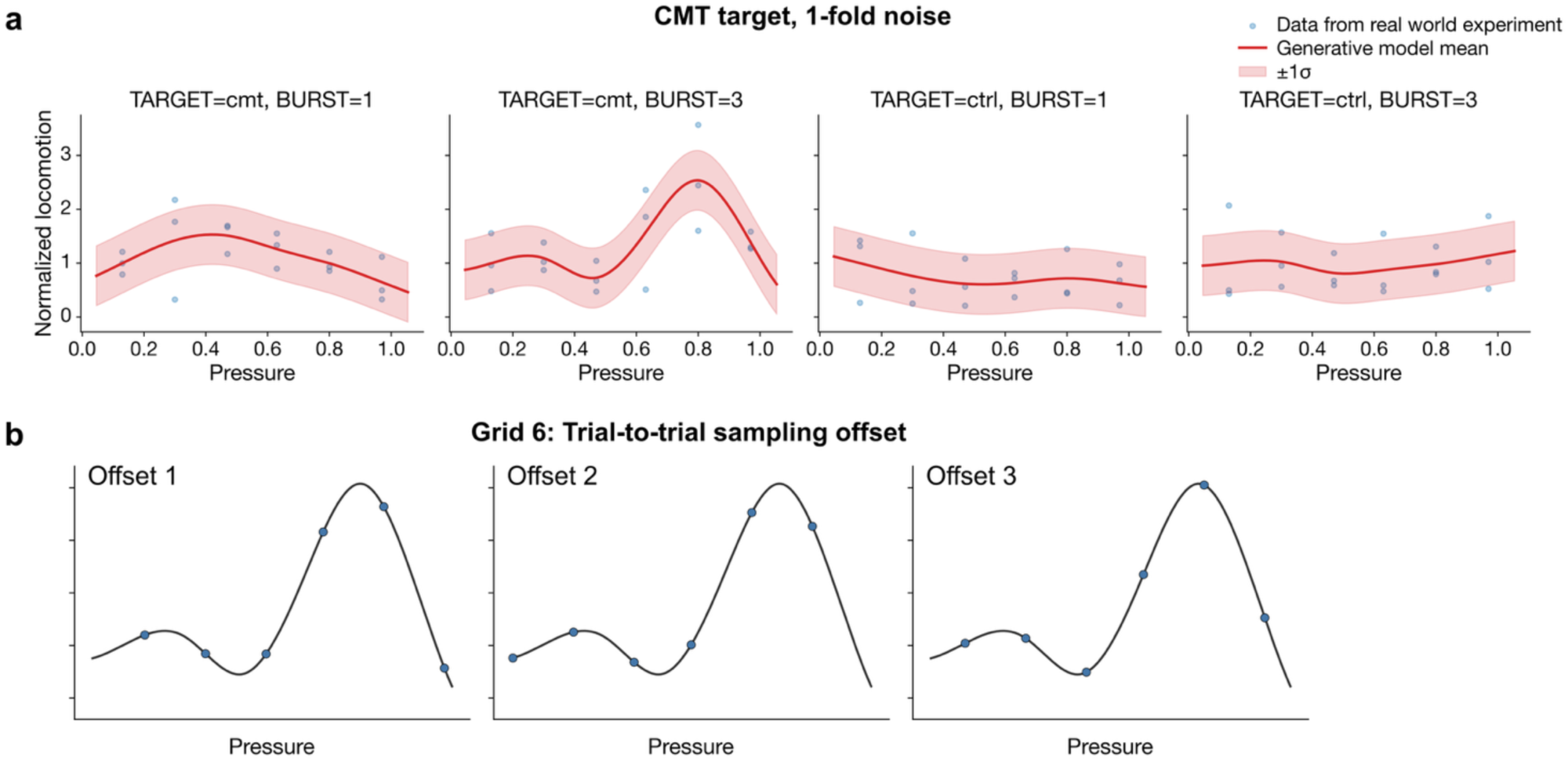
(**a**) Generative model for the *in-silico* experiments in the central medial thalamic (CMT) target with 1-fold noise standard deviation. (**b**) Representative trial-to-trial offsets introduced in the regular grid sampling to account for real-world uncertainties introduced by the a-priori definition of the grid. The plots show three examples for the Grid 6 configuration.

**Supplementary Figure 5.**
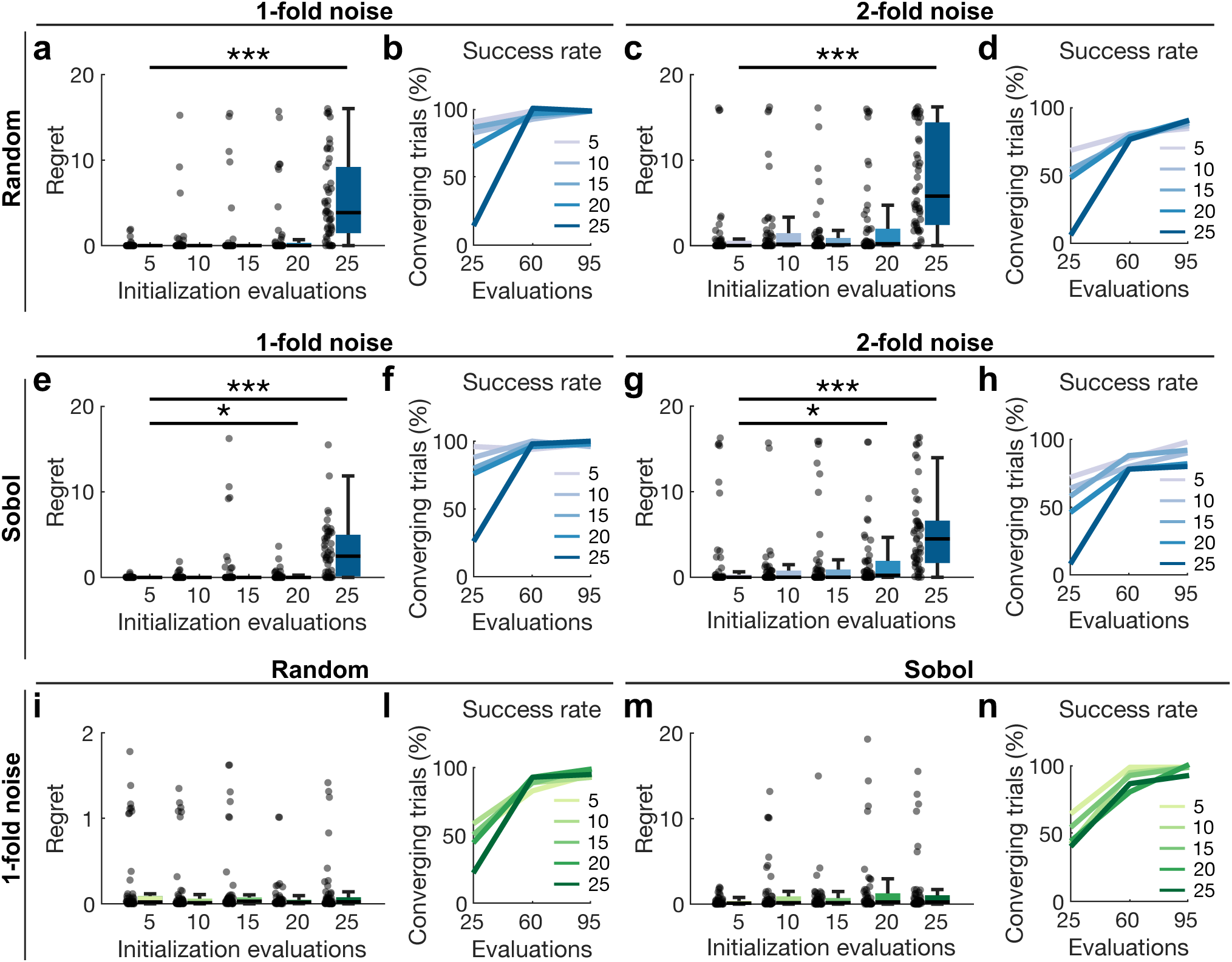
Effect of the initialization strategy on optimization metrics in the SC (**a-h**) and CMT (**i-n**) *in silico* experiments. (**a,e**) Regret at evaluation 25 with varying random- and Sobol-distributed initial points in the condition with 1-fold noise. We observed a significant effect for the number of initial points (*P* = 1.22 × 10^-36^ in a Kruskal-Wallis test). (**a**) Within the random initialization strategy, we observed a significant difference between 5 and 25 number of initial points; *P* = 1.38 × 10^-13^ (***) in B-H–corrected Mann-Whitney U-tests. (**e**) Within the Sobol initialization strategy, we observed a significant difference between 5 and 20 and 5 and 25 number of initial points; *P* = 0.03 (*) and *P* = 1.31 × 10^-11^ (***) in B-H–corrected Mann-Whitney U-tests. (**b,f**) Success rate (regret lower than 1% of the ground-truth minimum) for the condition with 1-fold noise and varying random and Sobol-distributed initial points. (**c,g**) Regret at evaluation 25 with varying random- and Sobol-distributed initial points in the condition with 2-fold noise. We observed a significant effect for the number of initial points (*P* = 7.41 × 10^-27^ in a Kruskal-Wallis test). (**c**) Within the random initialization strategy, we observed a significant difference between 5 and 25 number of initial points; *P* = 4.86 × 10^-11^ (***) in B-H–corrected Mann-Whitney U-tests. (**g**) Within the Sobol initialization strategy, we observed a significant difference between 5 and 20 and 5 and 25 number of initial points; *P* = 0.04 (*) and *P* = 1.96 × 10^-8^ (***) in B-H–corrected Mann-Whitney U-tests. (**d,h**) Success rate (regret lower than 1% of the ground-truth minimum) for the condition with 2-fold noise and varying random- and Sobol-distributed initial points. (**a,e** and **c,g**): In both the 1-fold and 2-fold noise conditions: there was no significant difference between random and Sobol sampling (*P* = 0.20 and *P* = 0.30 in a Kruskal-Wallis test, respectively). (**i,m**) Regret at evaluation 50 with varying random- and Sobol-distributed initial points in the condition with 1-fold noise. There was no significant effect of the number of initial points (*P* = 0.59 in a Kruskal-Wallis test). There was no significant difference between random and Sobol sampling (*P* = 0.58 in a Kruskal-Wallis test) (**l,n**) Success rate (regret lower than 5% of the ground-truth maximum) for the condition with 1-fold noise and varying random- and Sobol-distributed initial points.

**Supplementary Figure 6.**
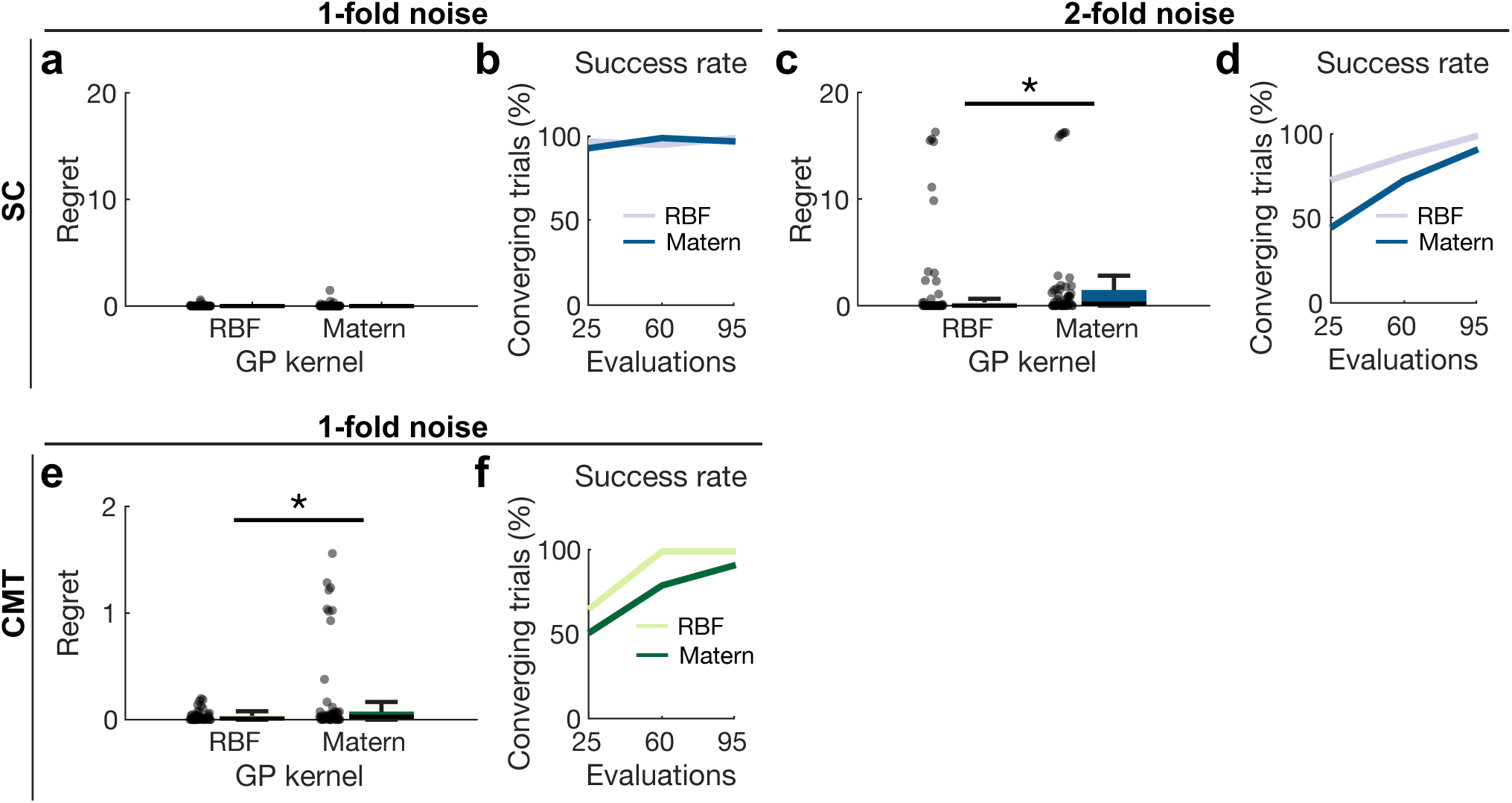
Effect of the GP kernel on optimization metrics in the SC (**a-d**) and CMT (**e,f**) in silico experiments. (**a**) Regret at evaluation 25 with varying RBF and Matérn GP kernels in the condition with 1-fold noise and 5 Sobol-distributed initial points. There was no significant difference between RBF and Matérn kernels; *P* = 0.72 in a Mann-Whitney U-test. When marginalizing across all numbers of initial points and both random and Sobol initialization strategies, there was no significant difference between RBF and Matérn kernels; *P* = 0.122 in a Mann-Whitney U-test. (**b**) Success rate (regret lower than 1% of the ground-truth minimum) for the condition with 1-fold noise and varying RBF and Matérn GP kernels. (**c**) Regret at evaluation 25 with varying RBF and Matérn GP kernels in the condition with 2-fold noise and 5 Sobol-distributed initial points. We observed a significant difference between RBF and Matérn kernels; *P* = 0.027 (*) in a Mann-Whitney U-test. However, when marginalizing across all numbers of initial points and both random and Sobol initialization strategies, there was no significant difference between RBF and Matérn kernels; *P* = 0.088 in a Mann-Whitney U-test. (**d**) Success rate (regret lower than 1% of the ground-truth minimum) for the condition with 2-fold noise and varying RBF and Matérn GP kernels. (**e**) Regret at evaluation 50 with varying RBF and Matérn GP kernels in the condition with 1-fold noise and 5 Sobol-distributed initial points. We observed a significant difference between RBF and Matérn kernels; *P* = 0.024 (*) in a Mann-Whitney U-test. When marginalizing across all numbers of initial points and both random and Sobol initialization strategies, RBF had lower median regret than Matérn; *P* = 0.013 in a Mann-Whitney U-test. (**f**) Success rate (regret lower than 5% of the ground-truth maximum) for the condition with 1-fold noise and varying RBF and Matérn GP kernels.

**Supplementary Figure 7.**
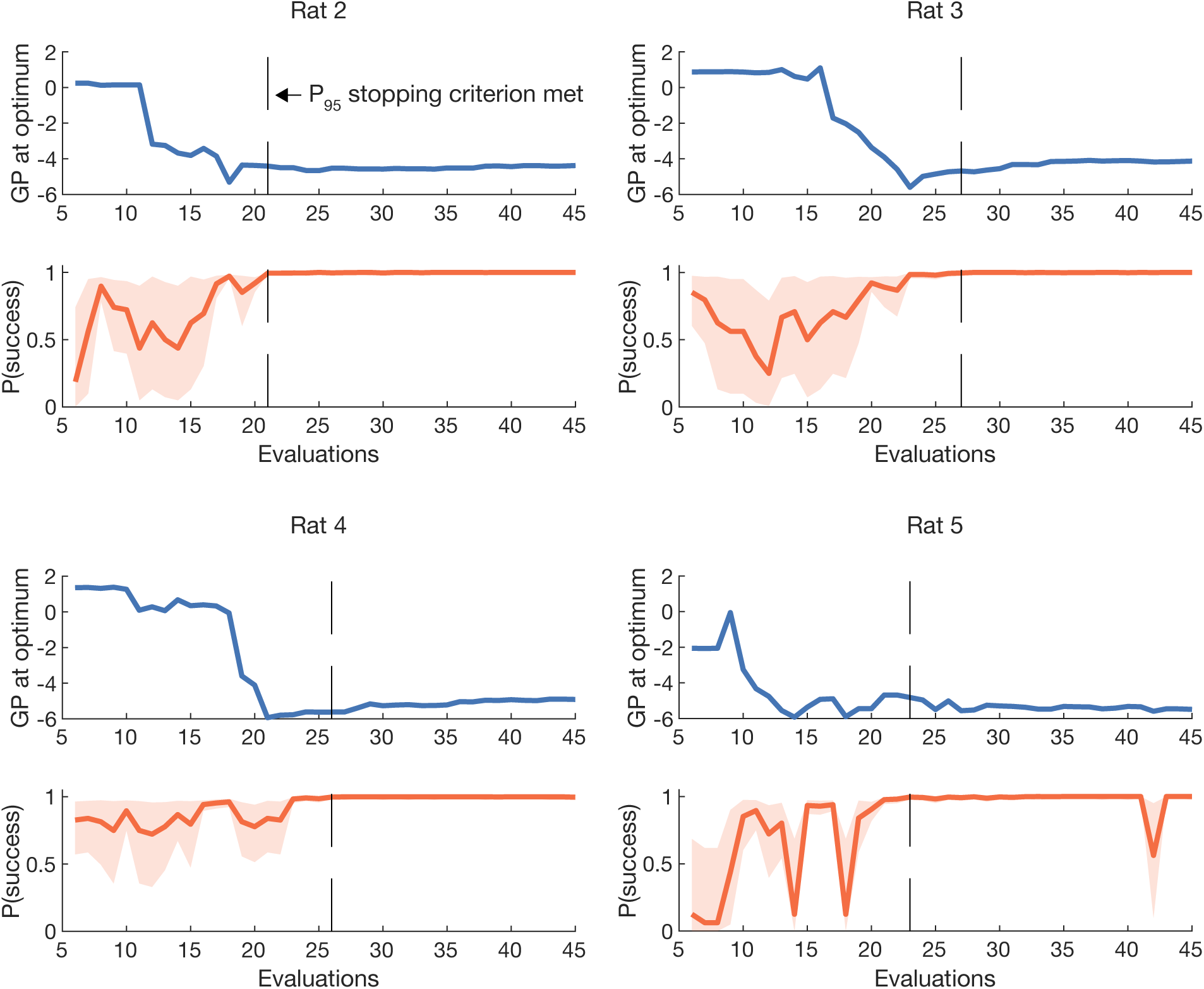
Time series showing the temporal evolution of the Gaussian process (GP) optimum and probability of success in all the live BEACUN experiments in the dorsal lateral geniculate nucleus. The orange shaded are represents the 95% confidence interval of the success probability. The dashed lines display the evaluation at which the P_95_ stopping criterion was met.

**Supplementary Video 1**: Evolution of the BECUN search and Gaussian process regression surrogate in a representative *in-silico* experiment targeting the centromedial thalamic (CMT) nucleus.

**Supplementary Videos 2, 3**: Evolution of the BEACUN search and Gaussian process regression surrogate for the live *in-vivo* experiments targeting the dorsal lateral geniculate (DLG) thalamic nucleus. Supplementary Video 2 shows the search for the rat displayed in Figure 5d. Supplementary Video 3 shows the search for Rat 4 in Supplementary Figure 7.

